# F_0_-F_1_ coupling and symmetry mismatch in ATP synthase resolved in every F_0_ rotation step

**DOI:** 10.1101/2021.11.14.468453

**Authors:** Shintaroh Kubo, Toru Niina, Shoji Takada

## Abstract

The F_0_F_1_ ATP synthase, essential for cellular energy production, is composed of the F_0_ and F_1_ rotary motors. While both F_0_ and F_1_ have pseudo-symmetric structures, their symmetries do not match. How the symmetry mismatch is solved remains elusive due to missing intermediate structures of rotational steps. Here, for ATP synthases with 3- and 10-fold symmetries in F_1_ and F_0_, respectively, we uncovered the mechanical couplings between F_0_ and F_1_ at every 36° rotation step via molecular dynamics simulations and comparison of cryo-electron microscopy structures from three species. We found that the frustration is shared by several elements. The F_1_ stator partially rotates relative to the F_0_ stator via elastic distortion of the b-subunits. The rotor can be distorted. The c-ring rotary angles can be deviated from symmetric ones. Additionally, the F_1_ motor may take non-canonical structures relieving stronger frustration. Together, we provide comprehensive understanding to solve the symmetry mismatch.

## Introduction

Adenosine triphosphate (ATP) is mainly synthesized by the enzyme F_0_F_1_ ATP synthase^1–3^. The enzyme is composed of two rotary motors, the F_0_ and F_1_ motor, and these motors are connected by two stalks, a central rotor and a peripheral stalk (Fig. 1a). The membrane-embedded F_0_ motor rotation driven by the proton motive force causes the central rotor rotation. The latter induces a series of conformational changes in the F_1_ motor, where ATP is synthesized from adenosine diphosphate (ADP) and inorganic phosphate (Pi). The F_0_F_1_ ATP synthase is also known as a reversible machine, which can hydrolyze ATP to pump protons against the proton motive force.

**Fig. 1.**
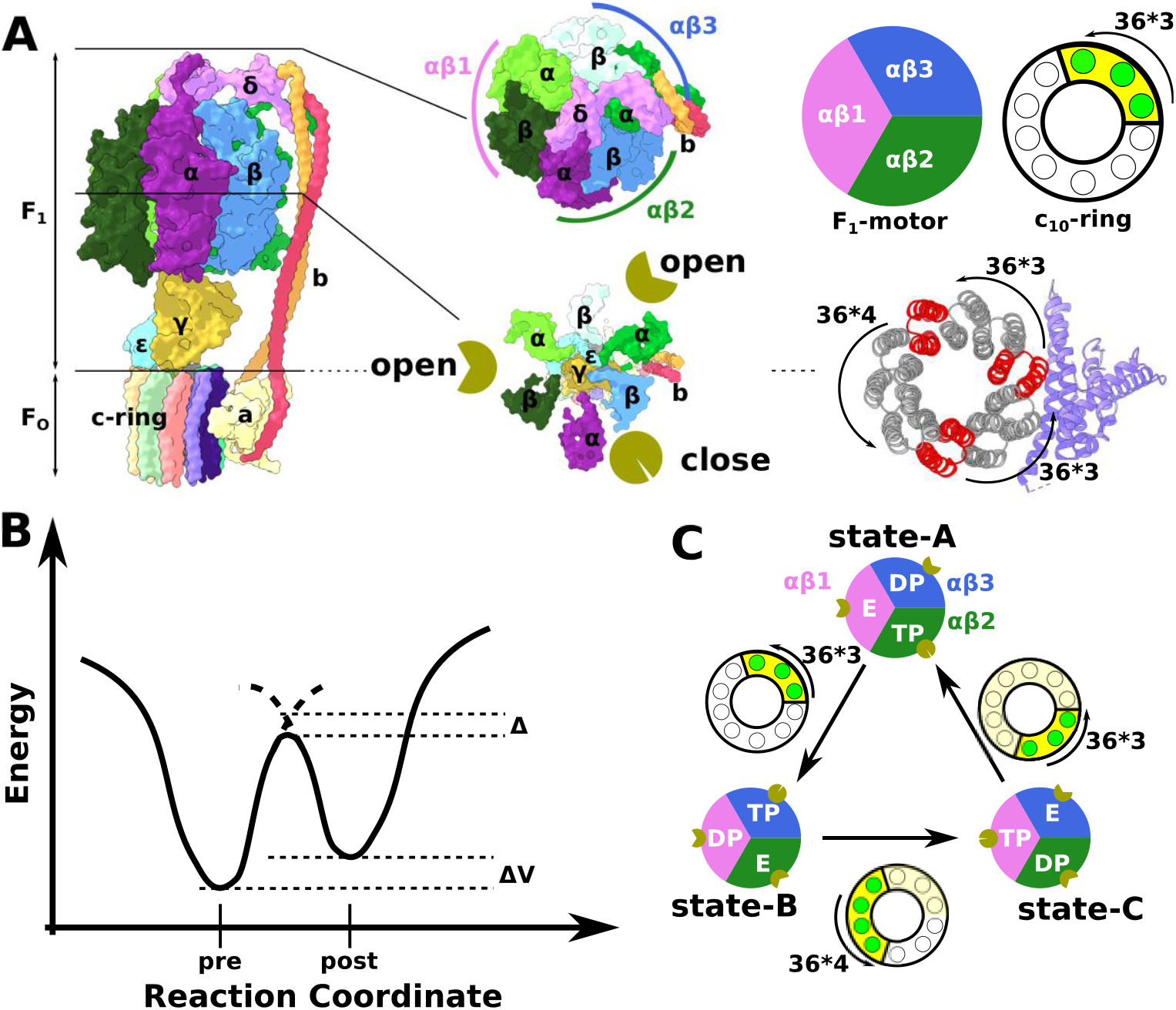
Structure of F_0_F_1_ ATP synthase and the simulation system. **a**. (Leftmost) The structure of *Bacillus* PS3 F_0_F_1_ ATP synthase holo-complex (the A state, PDB ID: 6N2Y). The F_1_ motor (top half) contains δ: pink; α_3_β_3_ hexamer made of three αβ pairs, αβ1: lime green and green; αβ2: violet and sky blue; αβ3: yellow green and white; γ: dark yellow; ε: cyan. The F_0_ motor contains c_10_-ring: pastel-colored barrel-shaped objects; a-subunit: pale yellow; b-subunits: pale red and orange. In the 6N2Y structure, αβ1, αβ2, and αβ3 take the E, TP, and DP states, respectively, and these take the open, closed, and open structures, respectively indicated by golden pie chart pictures. (Right) Guo *et al*. built three different ATP synthase conformations: A (PDB ID: 6N2Y), B (6N2Z), and C (6N30). The structure differences from A to B (AB), B to C (BC), and C to A (CA) contain three, four, and three 36°-c_10_-ring-positional-rotation in a counterclockwise direction, respectively. The red c-subunits indicate the ones closest to the a-subunit in the A, B, and C states. **b**. The schematic view of the double-basin model. “pre” and “post” indicate the minimum energy structures for the pre- and post-structures, respectively. *Δ* and *ΔV* are parameters to control the barrier height and relative stability between two minima. **c**. The conformational change cycle in the ATP synthesis mode.

The F_1_ motor contains the central rotor (γ/ε-subunit in bacterial F_1_) and an α_3_β_3_ hexamer made of three αβ dimers (Fig. 1a). Each αβ dimer has catalytic sites for the ATP synthesis reaction and take three distinct chemical states and conformations^4^, which are conventionally denoted as the ATP-bound state (TP for brevity), the ADP-bound state (DP), and the empty state (E). Especially, the β-subunit takes markedly different conformations among the three states; for the case of the *Bacillus* PS3 ATP synthase, which will be our target in this study, its C-terminal segment takes a “closed” conformation in the TP state, while it takes an “open” conformation in the DP and the E states^5^. Since a single-molecule measurement showed that the F_1_ motor in the ATP hydrolysis mode uncovered 120°-stepwise rotation of the central rotor, accompanied by progress of the chemical states in α_3_β_3_^6^. Thus, coupled with the nucleotide-dependent conformational change in the α/β-subunits, one round of the F_1_ motor results in the hydrolysis or synthesis of three ATP molecules, depending on the central rotor direction. The F_0_ motor is made of an a-subunit and a c-ring, which is a ring-shaped c-subunit oligomer working as a rotor (Fig. 1a). The number of c-subunits in the c-ring varies between 8-17 depending on the species^7–13^; 10 subunits in the *Bacillus* PS3 ATP synthase. Each c-subunit has one key proton-relaying residue. Single-molecule experiments for *Escherichia coli* ATP synthase (10 subunits) identified 36° stepwise c-ring rotations^14^. Thus, one round of the F_0_ motor was driven by the transfer of the same number of protons as that of c-subunits.

In contrast to the mechanisms of each motor, the mechanism of the coupling between the F_0_ and F_1_ motors is less clear. Because the number of c-subunits in the c-ring is not a multiple of three in most cases, the mean number of protons necessary for the synthesis of one ATP is not an integer value, which is sometimes called a symmetry mismatch. There has been much debate regarding the mechanism by which F_0_F_1_ ATP synthase resolves this mismatch^15^. For *Bacillus* PS3 ATP synthases, the two motors are connected via a central rotor made of c_10_γε and a peripheral stalk made of b_2_δ (Fig. 1a). Some early studies suggested the elastic distortion of the central rotor, γ-subunit^16–18^. Other studies have anticipated the role of δ^19^. Recent cryo-EM structures of mitochondria and *Bacillus* PS3 ATP synthases point to distortion in b-subunits^5,20^. From recent cryo-EM studies, the symmetric mismatch and the relatively rigid rotor together imply that the α_3_β_3_ hexamer rotate relative to the a-subunit^19,21^. Sobti *et al*. resolved three different rotational states and one sub-state in *E. coli* ATP synthase. The sub-state structure essentially had the same F_1_ configuration as one of the three state structures, but the c_10_-ring together with the F_1_ stator, c_10_α_3_β_3_γε, rotated relative to the F_0_ stator. Comparing individual conformations, we see that the b-subunits have significantly different conformations. Therefore, the existence of the sub-state and the flexibility of the b-subunit may be necessary for resolving the symmetry mismatch. Notably, however, the cryo-EM models in Sobti *et al*. include structural models for four c_10_-ring rotation states out of ten possible rotational states. Perhaps other states are more fragile and/or more short-lived, so that high-resolution models could not be built. Moreover, cryo-EM models provide static snapshots, as usual; for example, how the sub-state arises, how the b-subunit distortion resolves the mismatch, and when and how the F_1_-motor undergoes structural changes in response to the c-ring rotation remain unclear.

In this study, to address the F_0_-F_1_ coupling and the molecular mechanisms to solve the symmetry mismatch, we performed molecular dynamics (MD) simulations that mimic the ATP synthesis dynamics for one round of rotation in the holo-complex of the *Bacillus* F_0_F_1_ ATP synthesis. While many molecular simulations have been reported so far for each of the F_0_ and F_1_ motors^18,22–28^, to the best of our knowledge, this study provides MD simulations for the first time for one round of ATP synthase holo-complex. Our dynamic simulations showed the order and timing of the structural changes in the three *αβ*-pairs in the F_1_ motor during ten 36°-rotation steps of the F_0_ motor. Furthermore, relaxation simulations with a fixed c-ring rotation angle in every 36°-step exhibited the c-ring-dependent partial rotation of the F_1_ stator relative to the F_0_ stator and some twists in the rotor. Then, motivated by the simulation results, we conducted a comparative analysis of cryo-EM structures from three species, accounting for the structural changes in the ATP synthases. Together, we reveal how to solve the symmetry mismatch in F_0_F_1_ ATP synthesis in unprecedented detail.

## Results and Discussion

### Simulating the rotation dynamics in the ATP synthase holo-complex

The *Bacillus* PS3 ATP synthase cryo-EM study provides holo-complex structure models in the three rotational states: A (PDB ID: 6N2Y, Fig. 1a left), B (PDB ID: 6N2Z), and C (PDB ID: 6N30)^5^. In all of these holo-complex models, F_1_ was in the catalytic dwell. We introduce the numbering of the three αβ pairs, αβ1, αβ2, and αβ3, counterclockwise starting from the α/β-subunits furthest from the b-subunit (Fig. 1a, middle). In the A state, αβ1, αβ2, and αβ3 are in the E, TP, and DP states, respectively. The structural change from A to B (the AB process) involves the counterclockwise rotation of c_10_-ring by 3× 36°, together with the progress of the chemical steps in the three αβ pairs; αβ1, αβ2, and αβ3 change to the DP, E, and TP states, respectively. Similarly, the transitions from B to C (the BC process) and from C to A (the CA process) involve rotations by four and three 36° c-ring rotation steps, respectively, coupled with the chemical state changes in αβ pairs. We call this whole process the 3-4-3 pathway, assuming that the entire process starts and ends in the A state.

Given the 3-4-3 pathway in the three cryo-EM rotational states, we envisioned to simulate the process that the F_1_ motor in the A state would reach the B state when the F_0_ c_10_-ring rotates counterclockwise by 3×36°. To simulate the conformational change in the AB process, we need to model in a way that each αβ pair can accommodate both A and B state conformations. For this purpose, we employed a double-basin model that encodes A and B state conformations separately for each αβ system (see Methods for details). The double-basin model concisely builds two distinct conformational basins controlling both the energy barrier in between and the relative stabilities of the two basins^29–31^ (Fig. 1b). Also, for the BC and CA processes, introducing the double-basin models that connect the B and C states and that connect the C and A states, we expected to observe the respective state transitions in the F_1_ motor while the c_10_-ring rotates 4×36° and 3×36° degrees, respectively (Fig. 1c). To ensure the energetics of the ATP synthesis process, we set the summation of the relative stabilities of the three αβ double-basin models in the AB, BC, and CA processes to ∼ +36 kcal, which approximates the free energy increase for the synthesis of 3 ATP molecules from ADP and Pi.

In the simulations, we rotated the c_10_-ring by 36° with a constant angular velocity over 10^6^ MD steps, followed by a c_10_-ring pause for 9×10^6^ MD steps. Note that 10^4^ MD steps were denoted as one frame. For each case, we repeated the same simulations ten times with different stochastic forces. The structural transitions in each αβ are monitored by its reaction coordinate χ, which takes negative and positive values when αβ is in the pre- and post-states in the double-basin system, respectively. For the AB process, we repeated this rotation step four times (the upper c-ring panel of Fig. 2ab). For the BC and CA processes, we began with 10×10^6^ MD equilibration steps in the initial c_10_-ring angle and then repeated the rotation step four times (the upper c-ring panel of Fig. 2cd and Fig.2e,respectively).

**Fig. 2.**
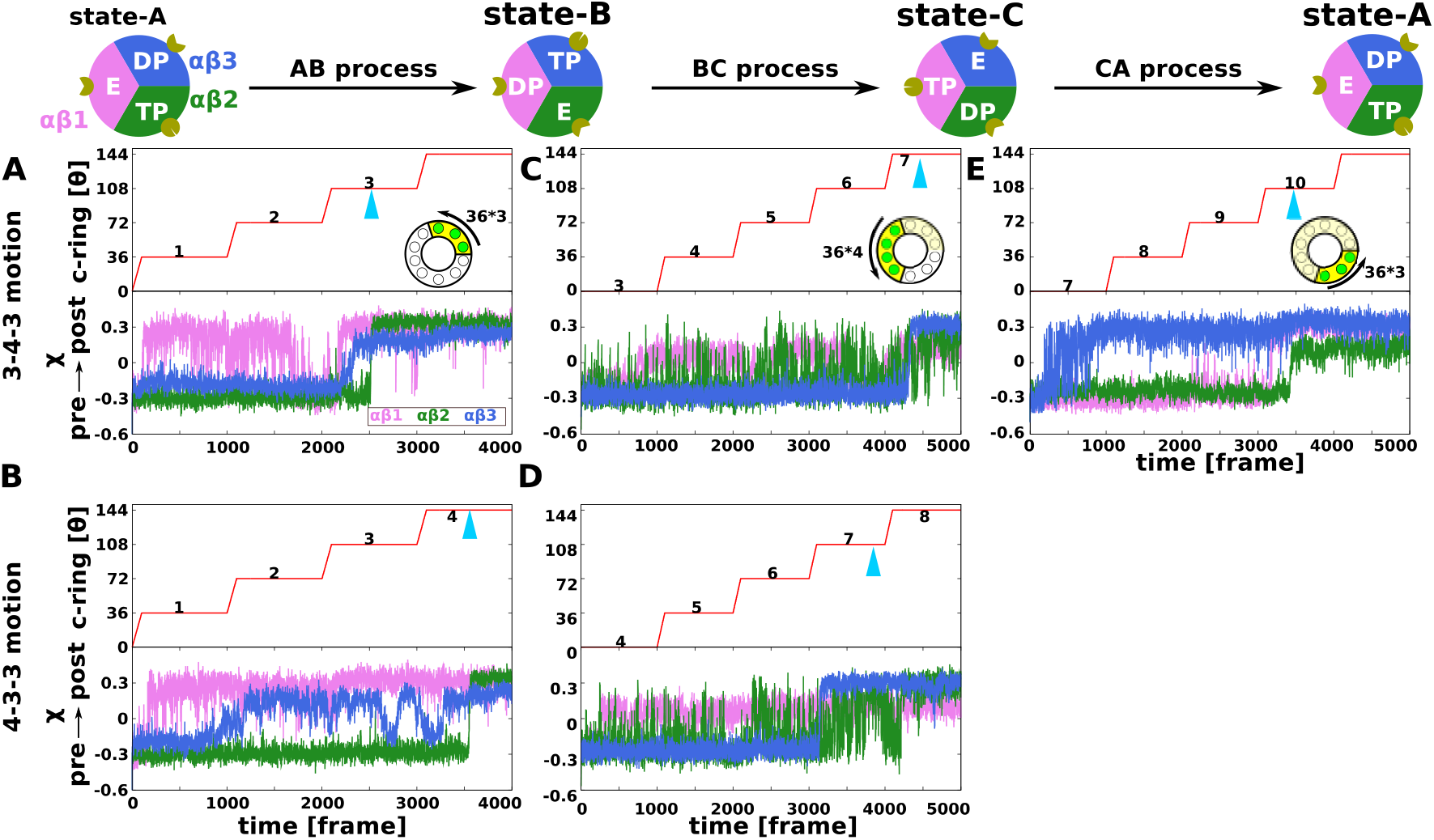
Structure change in the F_1_ motor during MD simulations of one round c_10_-ring rotations. **a-e**. Representative MD simulation trajectories for the AB (panel A), BC (panel C), and CA (panel E) processes in the 3-4-3 pathway, and for the AB (panel B) and BC (panel D) processes in the 4-3-3 pathway. Results of all the 10 trajectories are given in Supplementary Fig. 1. Each panel shows the c_10_-ring rotation time schedule (upper) and the reaction coordinate for the structure changes: χ of the αβ1, αβ2, and αβ3 (red, green, and blue, respectively) (bottom). The triangle mark colored cyan shows the timing when all the three αβ’s completed their structural changes. The CA process trajectories in the 4-3-3 motion is shown in Supplementary Fig. 2. One frame of time corresponds to 10^4^ MD steps.

### How many F_0_ c-ring rotation steps are necessary to induce the F_1_ nucleotide state transition?

We monitored the structural changes in each αβ (Fig. 2, χ pre → post) as well as other structures. When all three χ’s in F_1_ changed from negative to positive values, we regarded F_1_ to have completed its transition. Fig. 2a shows a representative trajectory of the AB process: (1) The αβ1 started transitions from the E state to the DP state at the ∼100th frame immediately after the c-ring rotated 36°, followed by rapid transitions back and forth. (2) The αβ3 made a transition from the DP to the TP state at the ∼2200th frame when the c-ring completed the third 36°rotation. (3) Finally, αβ2 transited from the TP to the E state at the ∼2500th frame, corresponding to the dissociation of a synthesized ATP molecule. At this stage, all three αβ pairs completed their transitions (blue triangle in the figure). This was within the third rotation step. Thus, in this trajectory, three c-ring rotation steps induced a complete structural transition in F_1_ during the AB process. Of the ten repeated trajectories, six showed similar results; three c-ring rotation steps induced complete F_1_ transitions. In the two trajectories, four c-ring rotation steps were necessary to induce F_1_ transitions (one case depicted in Fig. 2b). The remaining two did not complete the F_1_ transition at the end of the simulations (Supplementary Fig. 1a). Thus, as expected from the cryo-EM structures, we observed that in the majority of cases, the F_1_ structural transitions in the AB process were completed with the three 36°-c-ring-rotation steps.

Next, we describe the BC process. For this process, we first took the 3000th frame snapshot in the representative trajectory of the AB process (shown in Fig. 2a) as the initial structure. Fig. 2c shows the typical trajectory of the BC process. In this trajectory, the αβ1 and αβ2 pairs exhibited unstable fluctuations until αβ3 transitioned from the TP to the E state at the ∼4200th frame. Once αβ3 is in the E state, both αβ1 and αβ2 are stabilized in the TP and DP states. The overall structural change in F_1_ was completed after the fourth c_10_-ring rotation step. All trajectories which complete conformational changes (four out of ten) were completed with four c-ring 36°-rotation steps (Supplementary Fig. 1b). Therefore, we conclude that four 36°-rotation steps are necessary to induce complete structural changes in F_1_ in the BC process with this initial structure. We mentioned about the BC process on the 4-3-3 pathway in Supplementary Text 1.

Finally, the CA process is simulated. First, we started the simulation with the 5000th frame snapshot of the representative BC process trajectory in the 3-4-3 pathway. Fig. 2e shows a typical trajectory (Supplementary Fig 1d). Initially, αβ3 gradually changed from the E to the DP state between the 200th and 900th frames. Then, αβ1 made the transitions from the TP to the E state and settled down in the E state at the ∼ 3100th frame right when the third c-ring rotation step in the CA process was over. This is quickly followed by the transition in αβ2 from the DP to the TP state around the 3300th frame when the overall conformational change in this process was over. The F_1_ transition was completed within the third 36°-c-ring-rotation step in six of the ten trajectories tested. The F_1_ transition was completed within the second and fourth 36°-rotation steps, each in one trajectory. The remaining two trajectories did not exhibit complete structural changes. Therefore, the dominant pathway in the CA process completes the F_1_ state transitions by the three c-ring rotation steps. We mentioned about the CA process on the 4-3-3 pathway in Supplementary Text 1 and Fig. 2bd.

Altogether, our simulation predominantly showed the 3-4-3 pathway, which is in harmony with the cryo-EM studies. Additionally, we noticed a common feature as suggested in a classic work^1,32^: Of the three F_1_ αβ structure transitions in each process, the αβ that changes from the TP to the E state is a bottleneck that tends to decide the number of necessary 36°-rotation steps. ATP synthesis in this enzyme has the rate-limiting process of the product dissociation.

### Partial rotation of the F_1_ stator accompanied by the b-subunits bending

Recent cryo-EM studies by Murphy *et al*.^19^ and Sobti *et al*.^21^ reported more than three rotational states for ATP synthases from *Polytomella* sp. and *E. coli*, respectively, both of which contain a c_10_-ring. In both cases, they found varying degrees of rotation of the F_1_ stator relative to the F_0_ stator, raising the possibility of a partial rotation of the F_1_ stator as a mean to solve the symmetry mismatch. However, these structural models contain only a limited c-ring rotation step. How was the symmetry mismatch solved in every 36°-rotation step? To address this point, we calculated the rotary angle θ of the F_1_ stator relative to the F_0_ stator during the trajectories (Fig. 3a-e); here, the F_1_ stator and the F_0_ stator mean F_1_ α_3_β_3_ and F_0_ a-subunit, respectively. For this purpose, we fixed the rotation axis of the F_0_ c_10_- ring to the z-axis and F_0_ a-subunit to the positive x-axis and monitored the rotary angle θ of F_1_ α_3_β_3_ around the z-axis (further details in the Methods). Fig. 3a-e plot the time courses of θ for the same five trajectories as in Fig. 2a-e.

**Fig. 3.**
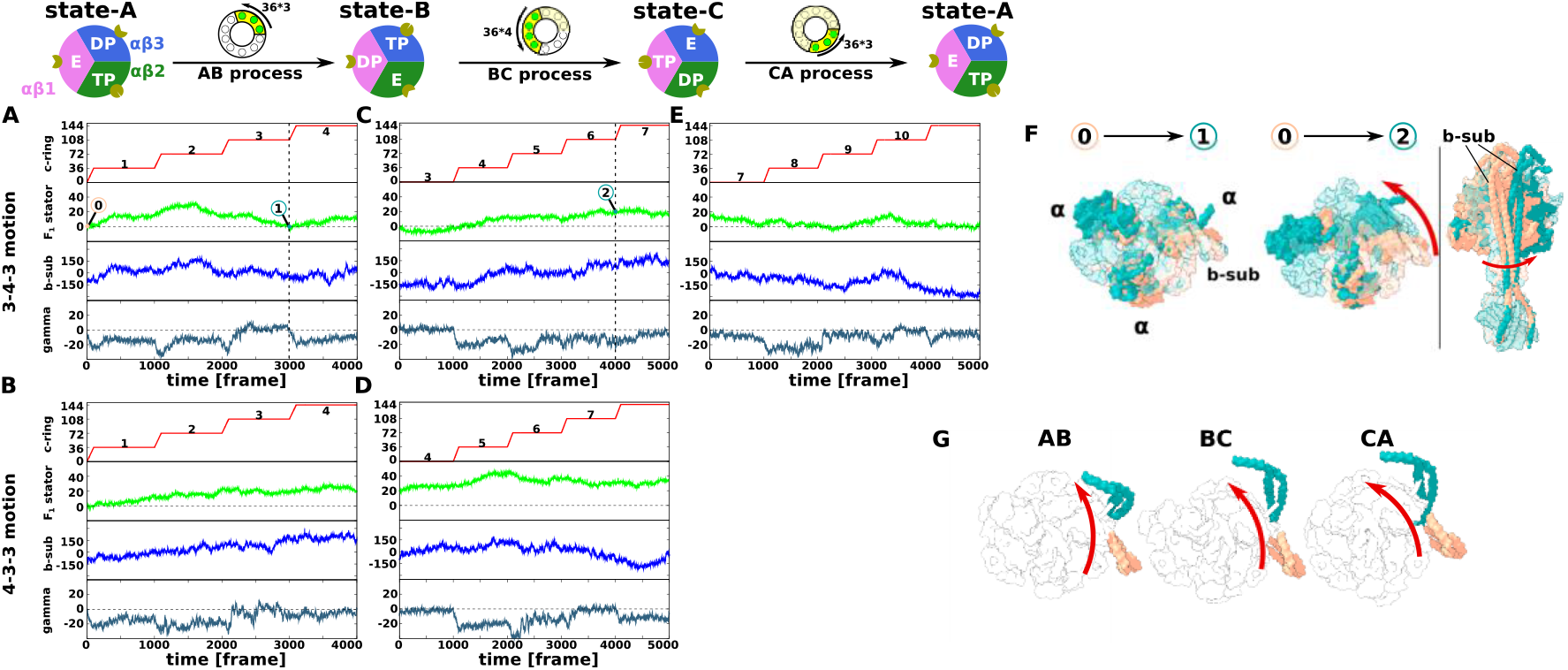
F_0_-F_1_ coupling during MD simulations of one round c_10_-ring rotations. **a-e**. Each panel plots, from the top to bottom, the rotation angle of the c_10_-ring, the rotation angle of the F_1_ stator, the first principal component about the b-subunit motions, and the rotation angle of the rotor defined by the angle of the upper part of γ-subunit relative to the rotation angle of c_10_-ring (the moving average over 10 frames) for the same trajectories as those shown in Fig. 2a-e. **f**. Superimposed snapshots shown in Fig. 3a-c trajectories 0, 1, and 2, respectively. The snapshot at time point 0 is colored dark salmon, and those at time point 1 and 2 are colored dark cyan. The three α-subunits are opaque to make it easier to see, and the others are translucent. **g**. The direction of structural change of the first principal component in the principal component analysis for the b-subunit. The first principal component value increases from dark salmon to dark cyan.

First, we examined the F_1_ stator angle θ of the AB process (Fig. 3ab). The F_1_ stator started to rotate ∼20° counterclockwise dragged by the c_10_γε rotation. In addition, the γ-subunit is distorted ∼-10° (the fourth panel of Fig. 3a). When the c-ring rotates counterclockwise, the rotation up to 10° tends to be absorbed by the distortion of the γ-subunit. In the subsequence process within the AB process in the 3-4-3 pathway (Fig. 3a), the F_1_ stator tends to return to the initial angle. Importantly, the return of the F_1_ stator angle occurred together with the structural transition in F_1_. Once F_1_ adopts conformations compatible with the cryo-EM structure of the B state, the F_1_ stator can accommodate the angle θ∼0 (Fig. 3f, left).

Next, we analyzed the simulation trajectories of the BC process in the 3-4-3 pathway (Fig. 3c). When the c_10_-ring started to rotate at the 1000th frame (the fourth cumulative c-ring rotation step is denoted as *n* = 4), the F_1_ stator is dragged by the rotor rotation, similarly to the above case (*n* =1). At the 1000th frame, while the c_10_-ring rotated by 36°, the F_1_ stator rotated by ∼20°. However, in sharp contrast to the above AB process, the F_1_ stator angle never returned to the initial angle (∼ 0°) until the simulation ended (*n* = 5 − 7) (Fig. 3f, right). At the end (the 5000th frame), the F_1_ stator was rotated by ∼20°. At this stage, the accumulated rotation in the c_10_-ring was 7×36°, counting from the A state, whereas the F_1_ motor made two progressive transitions rotating the chemical states by 2×120°. The difference in the rotation angle, 7×36°-2×120°=12°, may be adsorbed by the counterclockwise rotation of the F_1_ stator.

Notably, the rotation of the F_1_ stator in the BC process is accompanied by a large distortion of the b-subunit (the right-end extrusion in Fig. 3f right), in which a part that is close to F_1_ is rotated counterclockwise, whereas a part of the b-subunit bound to the F_0_ a-subunit remains in the original position. The distortion of the b-subunit is a passive conformational change. Owing to this distortion, elastic energy is charged into the b-subunit. We propose that the presence of the b-subunit, in addition to the balance of rotation angles discussed later, is the reason why the BC process requires more c-subunits rotation than other processes (Supplementary Text 2). To quantify the motion of the b-subunit, we performed principal component analysis (PCA) on the b-subunit; the first principal component (PC1) in the AB process represents a counterclockwise tilt along the F_1_ stator (Fig. 3g, left). The time course of the PC1 value (the third panel in Fig. 3a) indicates that it changes closely in parallel with the F_1_ stator angle θ. Note that a positive PC1 indicates a counterclockwise distortion of the b-subunit (the direction of the arrow in Fig. 3g).

Third, we examined the CA process of the 3-4-3 pathway. As shown in Fig. 3e panel, we observed a clockwise rotation of the F_1_ stator back to the initial angle. The b-subunit also followed clockwise tilting back to the relaxed structure in the A state (third panel in Fig. 3e). Therefore, it can be said that the CA process resolves the distortion caused by the counterclockwise rotation accumulated in the AB and BC processes. Lastly, we also mentioned the rotation of F_1_ stator for the 4-3-3 pathway in Supplementary Text 1.

### Relaxation simulations with the fixed c_10_-ring rotary angles

While these simulations provided dynamic view to resolve the symmetry mismatch, the obtained structures are inherently transient due to a relatively fast rotation of c_10_-ring. To obtain well-equilibrated structures at every 36°-c_10_-ring-rotation steps, which were not seen by cryo-EM studies, we conducted further MD simulations of relaxation. We fixed the c_10_-ring rotation step at *n* × 36 °-rotation steps for *n* =1 ∼ 10 and conducted further 2000-frame simulations ten times with the initial structure taken from the *n* × 1000th frame snapshots in the previous trajectories.

We found a clear trend in the AB process. For *n* = 1, we started the simulations from the 1000th frame snapshot of the representative trajectory (the red curve in Fig. 4a), as well as from the trajectory in which the F_1_ rotated the least amount at the 1000th frame among (the blue curve in Fig. 4a). Both sets of simulations steadily showed counterclockwise rotations of the F_1_ stator with varying degrees of rotation angles up to ∼30° (Fig. 4b, panel 1). Thus, our simulation models gave consistent results with recent cryo-EM studies^19^. We also monitored the distortion of rotor c_10_γε (Supplementary Fig. 3). In the *n* = 2 state, the F_1_ stator was rotated ∼20°, and then rotated clockwise to ∼ -10° in the c-ring state *n* = 3. Although the angle of the initial structure in *n* = 3 had a much more counterclockwise rotated, we found its clockwise rotation back to ∼0° during the relaxation simulation (the left-side green curve in Fig. 4a). Thus, the clockwise return of the F_1_ stator at *n* = 3 is a robust process.

**Fig. 4.**
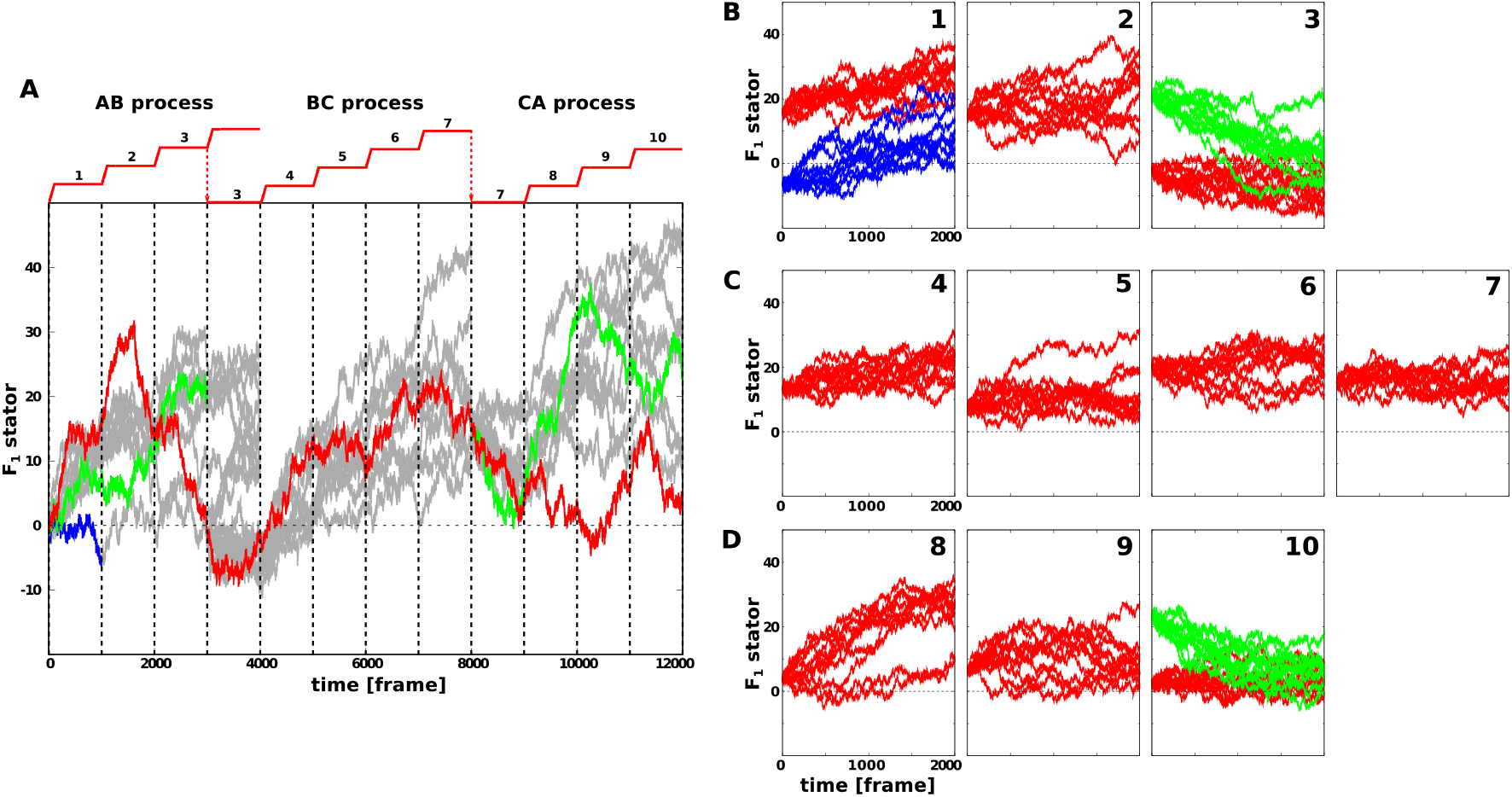
Relaxation simulation with the c_10_-ring rotation angle fixed at every n × 36° step. **a**. Reference trajectories from which relaxation simulations were conducted. The F_1_ stator rotation angle is depicted. Red: the representative one plotted in Figs. 2 and 3. Green in 0-3000th frame: one that was rotated markedly at the 3000th frame. Green in 8000-12000th frame: a case where the F_1_ stator was largely rotated at the end. Blue: a case where the F_1_ stator was least rotated. Gray: all the other trajectories. **b-d**. Each panel plots the time course of the F_1_ stator angle in the relaxation simulation with c_10_-ring fixed to the *n* × 36° rotation angle. Each simulation started from the snapshot of the reference trajectory in **a** at the corresponding time. **b**. the c_10_-ring rotation state *n* =1, 2, and 3 in the AB process. **c**. *n* =4, 5, 6, and 7 in the BC process. **d**. *n* = 8, 9, and 10 in the CA process.

In the BC process, we observed similar but not identical trends. In the c_10_-ring state *n* = 4, the F_1_ stator rotated counterclockwise by ∼20°, which is very similar to the c_10_-ring state *n* = 1. This angle was maintained in the c-ring states *n* = 5 and 6. Particularly, in the *n* = 6 state, we also observed marked, albeit incomplete, structural transitions in the F_1_ motor. Supplementary Fig. 4 shows that αβ3 exhibited irreversible and bottleneck transitions from the TP to E states in four of the ten trajectories. In the c-ring state *n* = 7, which is the final state in the BC process, the F_1_ stator is slightly, but not completely, turned to reach ∼+10°. A simple estimation may help our understanding: the c-ring rotated by 7×36° from the A state, while the F_1_ motor rotated by 2×120°. The difference was 12°.

Finally, in the CA process, we found the result to be somewhat similar to that of the AB process (Fig. 4d 8-10). In the c_10_-ring state *n* = 8, the F_1_ stator rotated counterclockwise by ∼30°, similar to the c_10_-ring state *n* = 1. We found that the rotor was significantly distorted at *n* = 8, which is different from the *n* = 1 case. The F_1_ stator slightly rotated back but remained rotated at ∼10° in the c-ring state *n* = 9. Finally, after the full round, the F_1_ stator settled down at the original 0°-angle. To check if the settled down was robustness, we simulated from another initial state in which the F_1_ stator rotated 20° (the right-side green curve in Fig. 4a). As we expected, we found the same settled down, therefore, we conclude that the F_1_ stator settles down to ∼0° when the system is fully relaxed after one round of the c_10_-ring.

In summary, we see that the F_1_ stator constantly rotates back and forth, dragged by the c_10_-ring rotation and by the chemical state change in F_1_, but eventually returns to its initial position after the full round. We have discussed the angle change from a typical trajectory, and in the next section, we will discuss it using more statistical values.

### On the symmetry mismatch between F_0_ and F_1_

The stepwise rotations of the F_0_ c_10_-ring and F_1_ motor shown in Fig. 5a (upper panel) illustrate the symmetry mismatch. By symmetry, the ideal elementary rotation steps of the c_10_-ring and F_1_ motor were 36° and 120°, respectively. We assume that the c_10_-ring and the F_1_ motor rotation steps coincided at the ground state angle, the A state, chosen as the angle 0° (this is an approximation, but it turned out reasonable). Then, the 120°-step of the F_1_ motor was flanked by 3×36 ° =108° and 4×36 ° =144° of the c_10_-ring steps with 108°-step closer. It is reasonable to assume that a smaller deviation in the angle corresponds to a lower energy of frustration. Thus, the 3×36 ° =108° step of the c_10_-ring may be realized to accommodate the F_1_ motor 120°-state. In the same way, the 240°-step of the F_1_ motor was flanked by 6×36 ° =216° and 7×36 ° =252° of the c_10_-ring steps with 252°-step closer. Obviously, the 360°-step was realized both by the c-ring and the F_1_ motor without frustration, which led to the 3-4-3 pathway, consistent with all three cryo-EM studies. This minimal-deviation-rule directly provides the simplest reasoning for the 3-4-3 pathway. The same rule may be applicable to c-rings composed of subunits other than ten (Fig. 5b). The c_8_-ring, the c_9_-ring, and the c_11_-ring may exhibit the 3-2-3, 3-3-3, and 4-3-4 pathways starting from the ground state.

**Fig. 5.**
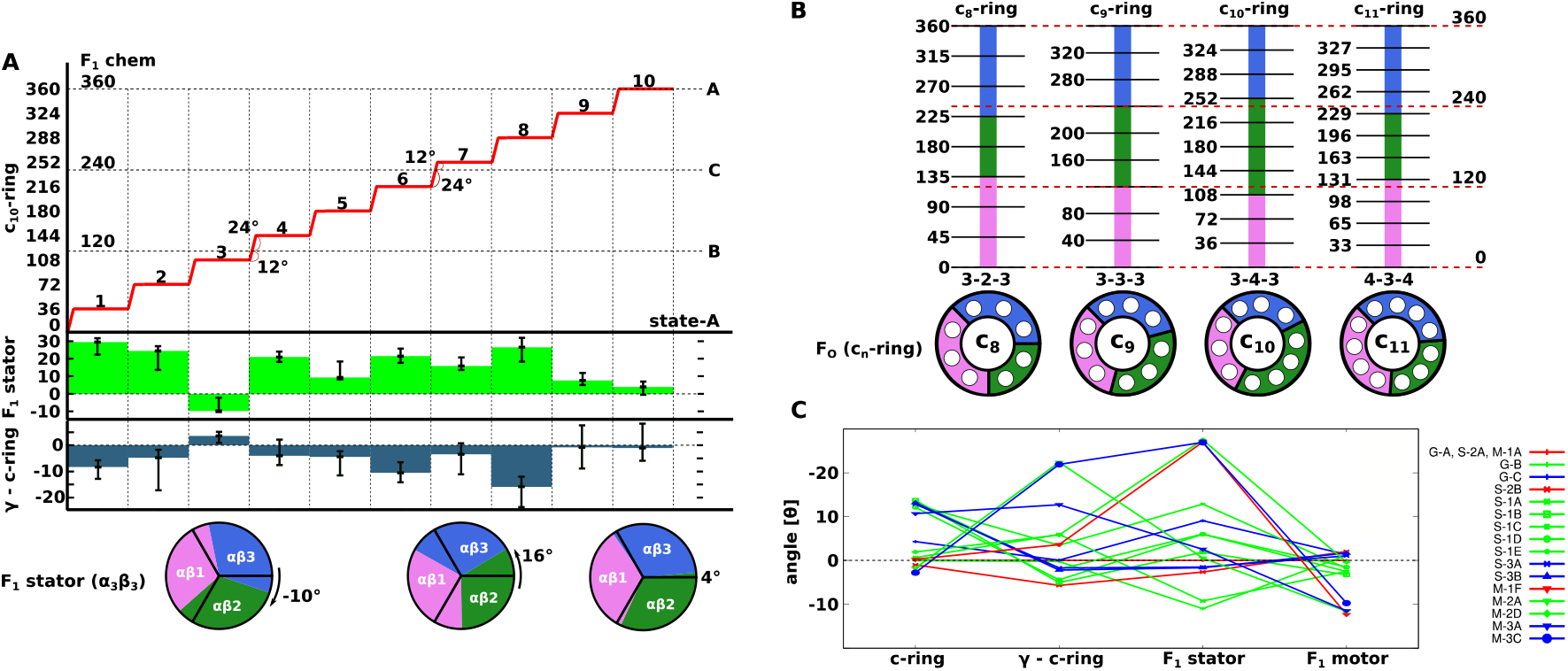
The F_0_-F_1_ coupling and the symmetry mismatch in ATP synthesis process. **a**. Summary of the symmetry mismatch and the elastic structure changes. The stepwise c_10_-ring rotation (the red ladder), the F_1_ motor rotation (the horizontal dashed line), the F_1_ stator rotation via the distortion of the b-subunit (the green bar in the second panel), and the rotor distortion (the blue bar in the third panel) are depicted for every n × 36° c_10_-ring rotation step. **b**. Predicted rotation pathways are described for systems with different number of c-subunits. **c**. Four elements of structure distortions found in cryo-EM studies. c-ring, the deviation of the c-ring rotation angle from its ideal angle. γ-c-ring, the rotary angle of the γ-subunit at its interaction site to αβ minus the c-ring rotation angle. F_1_ stator, the F_1_ stator rotation angle relative to the F_0_ stator. F_1_ motor, the rotation of the γ-subunit relative to the F_1_ stator α_3_β_3_.

The next question is how to resolve the mismatch between the F_0_ c_10_-ring and the F_1_ motor in the primary rotation states B (*n* = 3) and C (*n* = 7) in the 3-4-3 pathway, namely the difference between 120° and 108° in the B state and the difference between 240° and 252° in the C state. Structures provided by Guo *et al*. suggested that the mismatch of the range ∼12° can mostly be compensated by the rotation of the F_1_ stator via the distortion of the b-subunit (Table 1, Fig. 5c, and Supplementary Text 3).

**Table 1.** Rotary angles in cryo-EM structures of three ATP synthases.

| Guo | c<sub>10</sub>-ring | n<sup>1)</sup> | $\gamma - c_{10}$-ring<sup>2)</sup> | F<sub>1</sub> stator<sup>3)</sup> | F<sub>1</sub> motor<sup>4)</sup> | m<sup>5)</sup> |
| --- | --- | --- | --- | --- | --- | --- |
| 6N2Y (state-A) | 0 | 0/10 | 0 | 0 | 0 | 0/3 |
| 6N2Z (state-B) | 108.0(-0.0) | 3 | -0.2 | -11.0 | 118.8 (1.2) | 1 |
| 6N30 (state-C) | 247.7(4.3) | 7 | 0.1 | 9.1 | 238.7 (1.3) | 2 |
| Sobti |  |  |  |  |  |  |
| 6WNQ (2A) A<sup>Guo</sup> | 0 | 0/10 | 0 | 0 | 0 | 0/3 |
| 6OQV (2B) | -1.0(-1.0) | 0/10 | -5.7 | -2.6 | -2.0 (2.0) | 0/3 |
| 6OQR (1A) B<sup>Guo</sup> | 107.3 (0.7) | 3 | 6.0 | -9.2 | 122.5 (-2.5) | 1 |
| 6OQS (1B) | 130.3 (13.7) | 4 | -5.1 | 1.9 | 123.3 (-3.3) | 1 |
| 6OQT (1C) | 130.6 (13.4) | 4 | -2.0 | 6.0 | 122.5 (-2.5) | 1 |
| 6OQU (1D) | 131.9 (12.1) | 4 | -4.4 | 6.0 | 121.5 (-1.5) | 1 |
| 6PQV (1E) | 131.2 (12.8) | 4 | 3.5 | 12.9 | 121.8 (-1.8) | 1 |
| 6OQW (3A) C<sup>Guo</sup> | 239.1 (12.98) | 7 | -1.7 | -1.5 | 238.9 (1.1) | 2 |
| 6WNR(3B) | 238.8 (13.2) | 7 | -2.2 | -1.7 | 238.3 (1.7) | 2 |
| Murphy |  |  |  |  |  |  |
| 6RDH(1A) A<sup>Guo</sup> | 0 | 0/10 | 0 | 0 | 0 | 0/3 |
| 6RDW(1F) | 35.7 (0.3) | 1 | 3.6 | 27.1 | 12.3 (-12.3) | 0/3 |
| 6RDZ(2A) B<sup>Guo</sup> | 109.8 (-1.8) | 3 | 22.4 | 0.6 | 131.6 (-11.6) | 1 |
| 6RE8(2D) | 142.1 (1.9) | 4 | 5.8 | 27.5 | 120.4 (-0.4) | 1 |
| 6REB(3A) C<sup>Guo</sup> | 241.3 (10.7) | 7 | 12.7 | 2.6 | 251.5 (-11.5) | 2 |
| 6RES(3C) | 254.8 (-2.8) | 7 | 21.9 | 26.9 | 249.7 (-9.7) | 2 |

The angle value is in degrees. The value in parentheses is the deviation from the ideal value from its symmetry.
<sup>1)</sup> The c<sub>10</sub>-ring rotation step. <sup>2)</sup> The distortion of the rotor is defined by the difference between the angle of the upper part of the γ-subunit and the rotation angle of the c-ring. <sup>3)</sup> The rotation angle of the F<sub>1</sub> stator relative to the F<sub>0</sub> stator. <sup>4)</sup> The difference between the angles of the F<sub>1</sub> stator and F<sub>1</sub> rotor. <sup>5)</sup> F<sub>1</sub> motor rotation step.

The mismatch between the F_0_ c_10_-ring and the F_1_ motor increases after the 36° c-ring rotation from the primary states A, B, and C, namely the rotation states *n* =1,4, and 8. This range of mismatches cannot be absorbed by the F_1_ stator rotation. Instead, it is realized by the combinations of the three types of elastic structural changes in the structures of Sobti *et al*. and Murphy *et al*. (Fig. 5c and Supplementary Text 3): a) the b-subunits accommodate ±(7 − 14)°, which appears as the rotation of the F_1_ stator, b) the c_10_-ring rotation deviates ±(11 − 13)° from the ideal angles, and c) the rotor distortion is in the range of ±(4 − 11)°. In contrast, the F_1_ motor showed a fluctuation of only ∼±2°. It is a key feature that the F_1_ motor maintains the canonical angle with slight fluctuations in all cryo-EM structures. The corresponding angles monitored in our MD simulations are consistent with these experiments (Fig. 5a, the second and third panels), except that the MD simulations assumed uniform 36°-rotation steps of the c_10_-ring, which is not rigorously the case in cryo-EM structures.

In all the AB, BC, and CA processes, the first 36°-c-ring-rotation step from the primary rotation states induces the counterclockwise rotation of the F_1_ stator (*n* =1,4,8). This is in agreement with the findings of Murphy *et al*.^19^. The second 36°-c-ring-rotation step tends to keep the F_1_ stator rotated, which is often accompanied by significant distortion of the rotor and incomplete transitions in the F_1_ motor (*n* =2,5,9). The F_1_ motor may take structures different from canonical three-fold rotary states. These are strongly frustrated states and are likely too dynamic to be modeled at high resolution in cryo-EM studies. The simulations showed relatively diverse configurations. Here, we suggest an interesting scenario: In some of these states, the F_1_ motor may utilize the 80° sub-step to the so-called binding-dwell. Namely, the 80° sub-step of the F_1_ motor is close to 2×36°=72° step of c_10_-ring, the 200° sub-step is close to both 5×36°=180° and 6×36°=216° of c_10_-ring, and the 320° sub-step of the F_1_ motor is close to 9×36°=324° step of c_10_-ring, which may serve to relieve the highly frustrated energies. In the last 36°-c-ring-rotation step of the AB, BC, and CA processes, the F_1_ stator finally rotates clockwise back to near the original angle (*n* =3,7,10); and the final angles differ in the three processes: ∼-10°, ∼+10°, and ∼0° for the AB, BC, and CA processes, respectively.

## Conclusions

Using the recently obtained cryo-EM structures of the *Bacillus* PS3 F_0_F_1_ ATP synthases, we carried out MD simulations of the holo-complex that mimicked three cycles of ATP synthesis, the AB, BC, and CA processes, and one round of rotor rotation. We found that the AB, BC, and CA processes completed the respective structural changes in F_1_ with the highest probabilities when the c_10_-ring made three, four, and three 36°-rotation steps, which is consistent with the experimental results. At all ten 36°-step of c_10_-ring rotations, we investigated the holo-complex structural changes that resolve the symmetry mismatch between F_0_ and F_1_. The symmetry mismatch was resolved by the distortion of a few parts. First, the b-subunit distortion led to the rotation of the F_1_ stator back and forth relative to the F_0_ stator. Second, the rotor itself is distorted to a lesser extent. Third, the comparative analysis of cryo-EM structures from the three species showed that the c-ring rotary angles can be deviated from symmetric ones. Since the movement of the -subunit in αβ2 is suppressed by the b-subunit and *ε*-subunit, a stronger torque is required to overcome this barrier during the BC process. Simulation results, together with the comparative analysis of recent cryo-EM structure models, reveal molecular reasoning to resolve the symmetry mismatch.

Since we adopted a simple approach to simulates rotary motions of the ATP synthase holo-complex, it has some limitations. First, since we used a classical MD with a coarse-grained molecular representation, the chemical reaction itself was only dealt with conformational changes correlated with chemical reactions. Second, since we adopted a predefined time course of c_10_-ring rotation, we have not been able to reproduce the effects of stochastic proton transport. Third, since some parts of the b-subunit around δ-subunit was missing in cryo-EM models, we cannot deny the possibility that this missing region contribute to the conformational change. Despite these limitations, our MD simulations revealed the mechanical coupling between F_1_ and F_0_ with reasonable accuracy in detail, which is sufficient to fully explain the solution of the symmetry mismatch in ATP synthases.

## Materials & Methods

### Model building

We used the *Bacillus* PS3 ATP synthase structure model for the A state (Protein Data Bank (PDB) ID, 6N2Y) obtained by cryo-EM as the reference structure for the corresponding A state in our MD simulations^5^. The model consists of eight proteins and 22 subunits, ab_2_c_10_α_3_β_3_γδε. Based on the PDB structure, we modeled loops for the missing regions in the a-subunit using MODELLER^33^. For the sake of structural symmetry, the range of residues used for three α- and β-subunits was made the same: I8 to S501 for α-subunits and T2 to M469 for β-subunits. In addition, one of the b-subunits (called b2 in the original PDB data) contains a C-terminal region that is neither modeled nor has a sequence assigned. This region was excluded from the simulation system. We established the coordinate system such that the center of masses of the F_0_ c_10_-ring was set at its original point, the rotation axis of the c_10_-ring coincided with the z-axis, and R169 of the F_0_ a-subunit was on the x-axis.

We also prepared reference structure information for B and C states. Notably, we did not use the structural models given in the PDB for these states. Instead, we mostly used the structure information for the A state because we assumed that the A state was the ground-state form. In the structure-based simulation, the reference structure for each subunit was used to define the lowest energy state of the subunit, except for the structures of α_3_β_3_. Assuming a change in the chemical state and the corresponding change in the stable conformation, we repositioned the reference structures in a rotary manner. For example, the reference structure information of αβ1 in the B state, which is in the DP state, was copied from the reference structure of αβ3 in the A state, having the same DP state. Similarly, all reference structure information could be obtained from those in the A state.

### MD simulation setting

We performed coarse-grained MD simulations using the reference structure described above. In the coarse-grained representation, one amino acid was treated as one particle located at the Cα position. We primarily used the energy function AICG2+^34,35^, which has been intensively used in simulations of large molecular complexes^25,36,37^. In this function, the reference structure was assumed to be the most stable conformation, and many parameters inside were determined from the atomic interactions in the all-atom reference structures (detail in Supplementary Text 4).

In this study, to investigate the mechanical coupling between F_0_ and F_1_ parts, we designed a minimal setup that mimicked ATP synthesis reactions induced by the c-ring rotation driven by the proton-motive-force in a simple design. Assuming the c-ring rotation driven by the proton-motive force, we made the c-ring rotate with a predefined time course. In the F_1_ part, for each αβ pair that sandwiches the ATP catalytic site, we set multiple basins that encode ATP-bound (TP), ADP-bound (DP), and empty (E) state conformations. When the state transition settled to the new state, we considered that the corresponding chemical event occurred (detail in Supplementary Text 5).

In our minimal design, the F_0_ a-subunit was a rigid part of the F_0_ stator and was fixed to the initial structure and position. The F_0_ c_10_-ring was treated as a rigid body and rotated by design around the z-axis with a predefined schedule.

With these setups, we ran simulations on several processes: AB, BC, and CA. First, in the AB process, the simulation started from the reference structure built from PDB ID: 6N2Y. In the simulation, the c_10_-ring rotated stepwise 36° by 36° as designed. The coupling between F_0_ and F_1_ led the three double-basin systems of the F_1_-motor to exhibit their conformations and reach the post-state (state B). Then, we selected a representative snapshot from the trajectory and treated it as the initial model for the subsequent simulation of the BC process. The simulation of the BC process began with this model. After three or four c-ring 36°-rotation steps, we also picked a representative snapshot of the initial structure of the CA process. Finally, we performed a CA simulation. In each simulation, we repeated 10 MD runs with different stochastic forces using CafeMol version 2.1^38^. Unless otherwise noted, we took 4×10^7^ MD steps, 5×10^7^ MD steps, and 5×10^7^ MD steps for the AB, BC, and CA processes, respectively. We used underdamped Langevin dynamics at 323 K temperature and set the friction coefficient to 2.0 (CafeMol unit); default values were used for the others.

## Data Availability

The cryo-EM structures used in this paper are available download from the Protein Data Bank under PDB IDs 6N2Y, 6N2Z, and 6N30 for the *Bacillus* PS3 ATP synthase; 6WNQ, 6OQV, 6OQR, 6OQS, 6OQT, 6OQU, 6PQV, 6OQW, and 6WNR for the *E. coli* ATP synthase; 6RDH, 6RDW, 6RDZ, 6RE8, 6REB, and 6RES for the *Polytomella* sp. ATP synthase.

## Code Availability

All MD simulations in this paper was performed by CafeMol software. It can be downloaded from https://www.cafemol.org.

## Acknowledgements

S.K. was supported by JSPS Research Fellowship. This work was also supported by the MEXT grant JPMXP1020200101 as “Program for Promoting Researches on the Supercomputer Fugaku” (ST), partly by JSPS KAKENHI grants 20H0593 (ST) and 21H02441 (ST), and by by the Japan Science and Technology Agency (JST) grant JPMJCR1762 (ST).

## Author Contributions

S.K. and S.T. conceived and designed the project; T.N. developed the simulation code; S.K. performed the simulations; S.K. and T.N. analyzed the data; S.K. assembled figures, and all authors discussed the results and were involved in the manuscript writing process.

