## Supporting Figure, Text, and Table for "F_0_-F_1_ coupling and symmetry mismatch in ATP synthase resolved in every F_0_ rotation step"

### Supplementary Information

#### Supplementary Figures

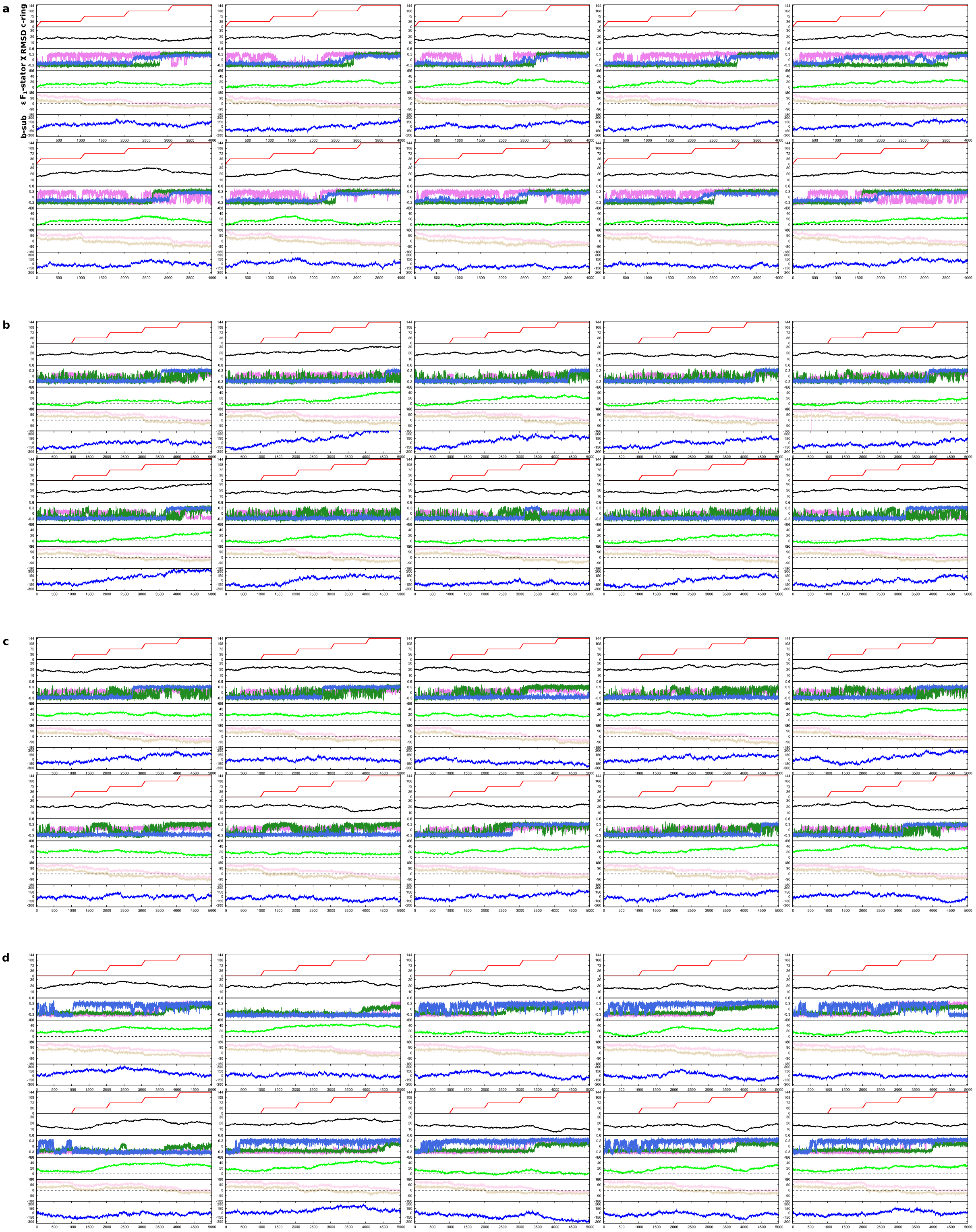

**Supplementary Fig. 1 All the MD simulation trajectories.**

All 10 MD simulation trajectories represent the AB, BC, and CA processes. Each panel shows, from the top to the bottom, the $c_{10}$-ring rotation angle (red), the RMSD from PDB ID 6N2Z, 6N30, and 6N2Y in the AB, BC, and CA processes, respectively (black), the reaction coordinates χ of αβ1, αβ2, and αβ3 (red, green, and blue, respectively), $F_1$ stator rotation angle (green), angle between ε- and α/β-subunit of system-2 (pale orange and pale pink, respectively), and the first principal component of the b-subunits (blue). **a**. The AB process. **b**. BC process in the 3-4-3 pathway. **c**. BC process in the 4-3-3 pathway. **d**. CA process in the 3-4-3 pathway.

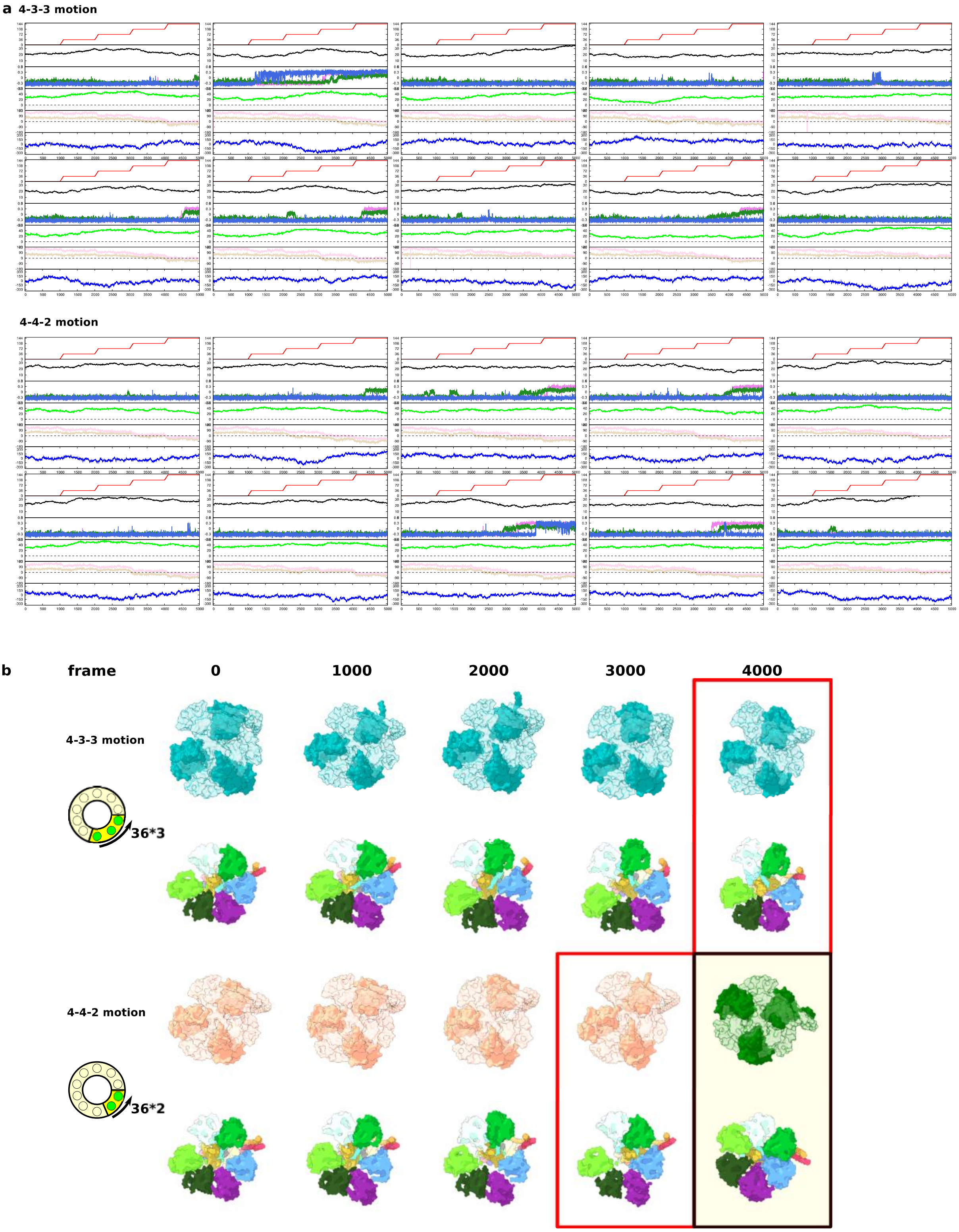


13

14 **Supplementary Fig. 2 CA process in the 4-3-3 and 4-4-2 pathways.**

15 **a.** The CA process trajectories in 4-3-3 and 4-2-2 pathways. Each panel shows the same quantities as

16 Supplementary Fig. 1. The above and bottom trajectories start from the 3800th and 4500th frames of

17 the BC process trajectory shown in the Fig. 2A lower and middle, respectively, thus corresponding to

the 4-3-3 and 4-2-2 pathways. **b.** Snapshots of every 1000 frames taken from each representative trajectory in Supplementary Fig. 2a. Three α-subunits are in dark cyan (for the 4-3-3 pathway) and dark salmon (the 4-4-2). The color-coded below are the section cut at the 256th residue of the three α-subunits. The two red boxes show the structures at the moment when the $c_{10}$-ring has just come to the initial position in each system. The black box is the original state-A structure shown in the same way with the other snapshots.

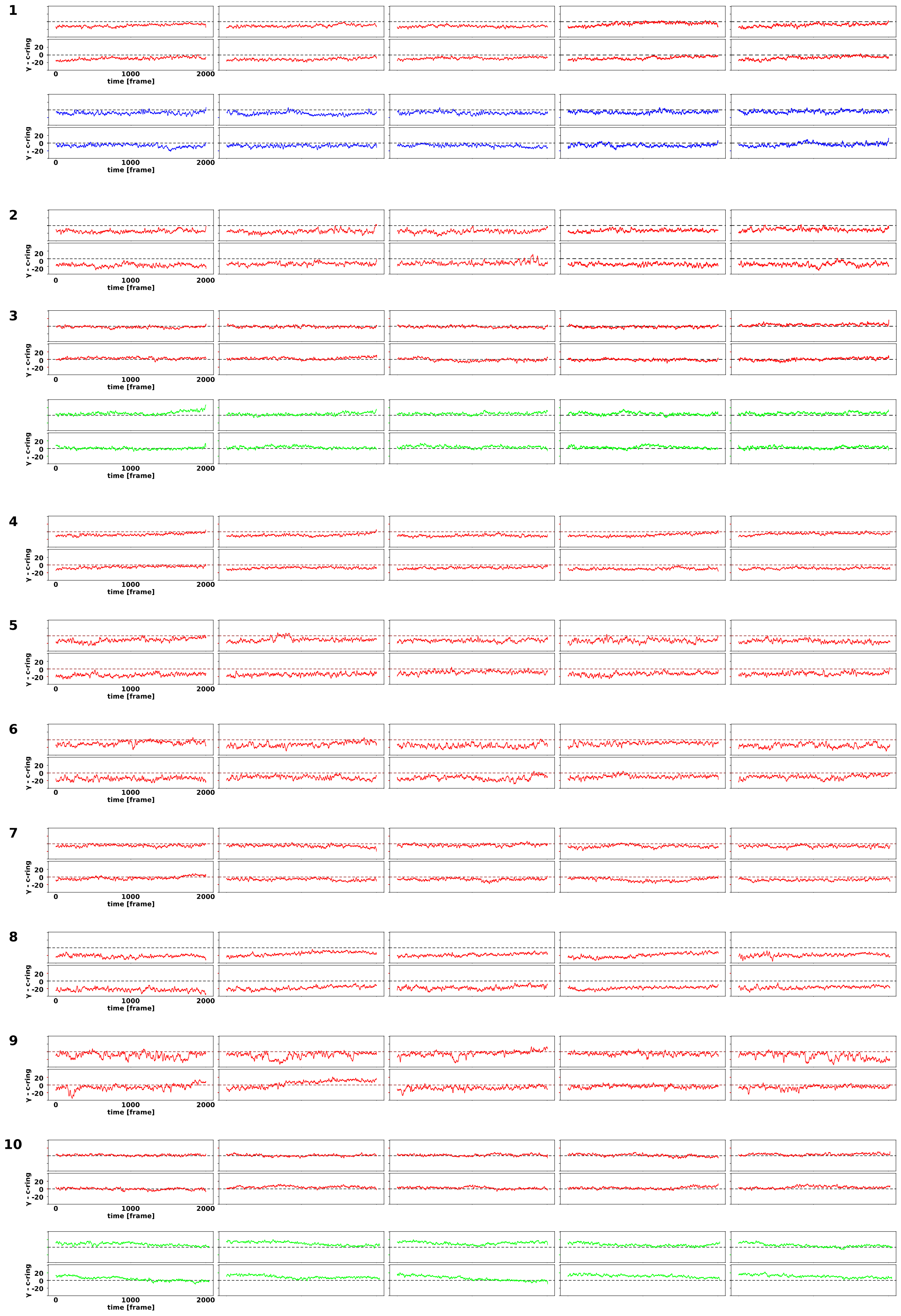


**Supplementary Fig. 3** **The rotor distortion trajectories in the relaxation simulations**

The rotor distortion trajectories for the relaxation simulations corresponding to Fig. 4bcd.

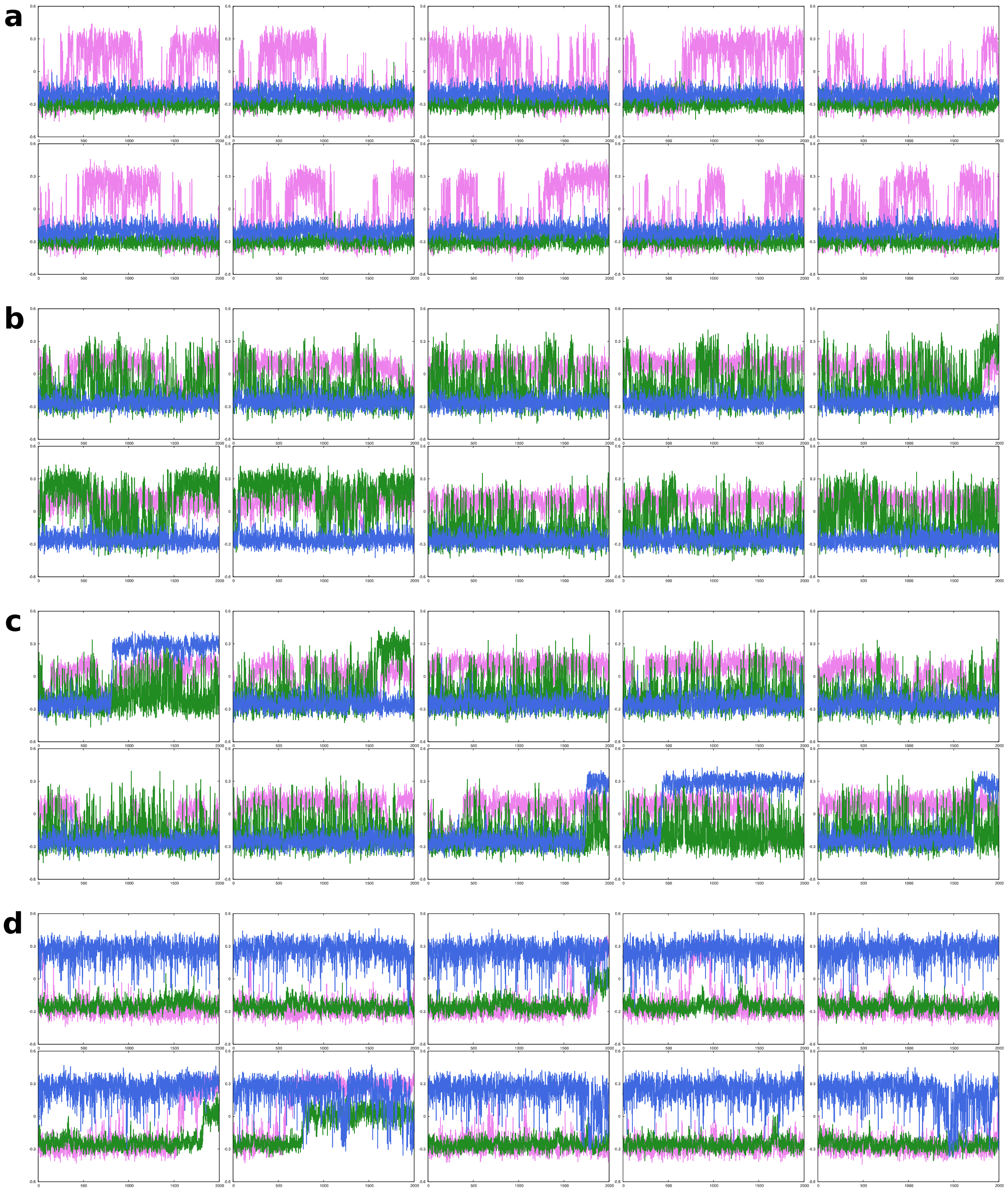

**Supplementary Fig. 4 The structure changes in αβ pairs in the relaxation simulations.**

The reaction coordinates for the structural changes are χ of αβ1, αβ2, and αβ3 (purple, green, and blue, respectively) in the relaxation simulations. **a.** $n = 2$ (Fig. 4B). **b.** $n = 5$ (Fig. 4C). **c.** $n = 6$ (Fig. 4C). **d.** $n = 9$ (Fig. 4D).

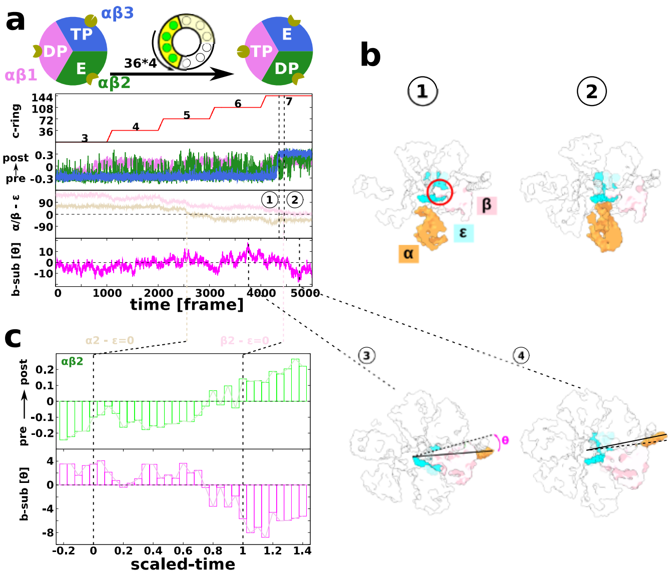


**Supplementary Fig. 5 Structural changes in αβ2 are controlled by ε- and b-subunits.**

**a.** A representative trajectory in the BC process (the same as Figs. 2C and 3C). From top to bottom: the rotation angle of the $c_{10}$-ring (red); the reaction coordinates $\chi$ to monitor the pre-to-post transition for αβ1, αβ2, and αβ3 in red, green, and blue, respectively; the angle ∠αOε and ∠βOε in pale orange and pale pink, respectively; and the angle ∠bOβ in magenta. **b.** Snapshots corresponding to time points marked by encircled numbers 1-4 in Fig. 5A, 1 and 2, where the α- and β-subunits of αβ2 are in orange and pink, respectively. In 3 and 4, β-, ε-, and b-subunits of αβ2 are in pink, cyan, and orange, respectively. **c.** The averaged changes of αβ2 reaction coordinate $\chi$ and angle ∠bOβ for β-subunit of αβ2 along the scaled time.

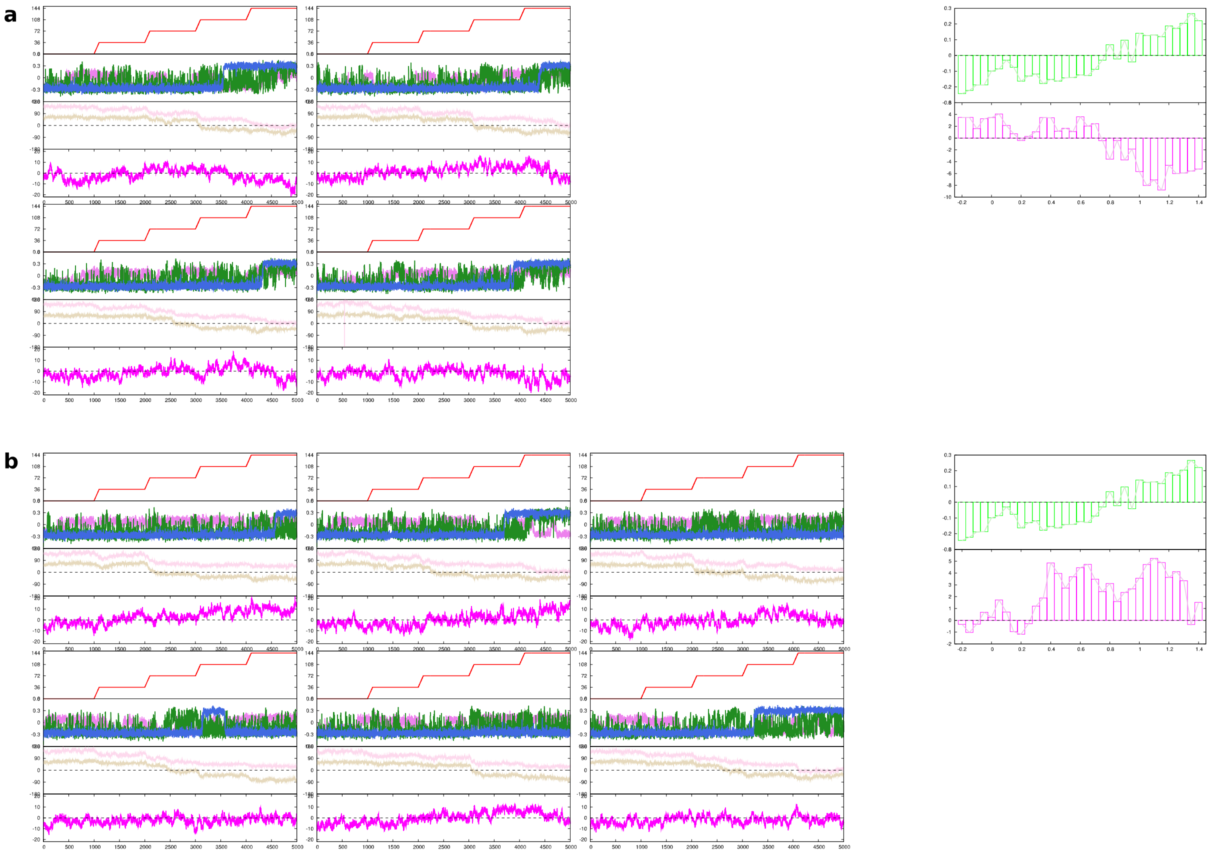

**Supplementary Fig. 6 The structural changes in αβ pairs and the angle between ε- and α/β-subunit of αβ2 on the scaled time.**

**a (Left).** The BC process trajectories in which all αs completed their conformation changes within the simulation time. **a (Right).** The changes in αβ2 χ-value and the angle between the b-subunit and the β-subunit of αβ2 on the scaled time axis were average from Supplementary Fig. 5a trajectories (the upper green and lower magenta, respectively) (same with Supplementary Fig. 5c). **b (Left).** The BC process trajectories in which some of the αs did not complete the structural change. **b (Right).** The changes in αβ2 χ-value and the angle between the b-subunit and the β-subunit of αβ2 on the scaled time axis were average from Supplementary Fig. 6b trajectories.

#### Supplementary Tables

**Supplementary Table 1** **The list of $\Delta$ and $\Delta V$.**

| | AB | | BC | | CA | | sum of $\Delta V$ |
|---|---|---|---|---|---|---|---|
| | $\Delta$[1)] | $\Delta V$ | $\Delta$[1)] | $\Delta V$ | $\Delta$[1)] | $\Delta V$ | |
| αβ1 | 600 | -60 | 860 | +5 | 770 | +80 | +25 |
| αβ2 | 770 | +40 | 600 | -45 | 860 | +23 | +13 |
| αβ3 | 860 | -85 | 770 | +55 | 600 | +22 | -3 |
| sum of $\Delta V$ | | -105 | | +15 | | +125 | +35[2)] |

[1)] The unit of $\Delta$ and $\Delta V$ is kcal/mol. $\Delta = 600\mathrm{kcal/mol}$ corresponds to the transition from E to DP, $\Delta = 770\mathrm{kcal/mol}$ corresponds to the transition from TP to E, and $\Delta = 860\mathrm{kcal/mol}$ corresponds to the transition from DP to TP.

[2)] Note that $\Delta V$ represents the shift in the internal energy of the post-state relative to the pre-state. The free energy difference should be different because of the difference in the conformational entropy. However, the sum of all the $\Delta V$ represents the free energy change in the $F_1$ motor during one round of ATP synthase rotation because the initial and final states should have the same conformational entropy.

#### Supplementary Movie.

**Supplementary Movie 1 Overview of ATP synthesis.**

The movie shows the results of the representative trajectory that achieved the 3-4-3 pathway (corresponding to the red curve in Fig. 4) for the AB (15 s), BC (25 s), and CA (20 s) processes. Top-left, the canonical view, which is the same view as Fig. 1a; bottom-left, 90° rotation from the canonical view around the z-axis; top-right, 90° rotation from the canonical view around the x-axis; bottom-right, a cross-section from the top-right view, which helps recognize the conformational changes. The color scheme is the same as that in Fig. 1a, except that the red color is used to highlight the C-terminal region of three β-subunits that cause significant nucleotide-dependent conformational changes.

#### Supplementary Text.

#### Supplementary Text 1

##### The 4-3-3 pathway.

In the two of ten trajectories in the AB process, four $c_{10}$-ring rotation steps were necessary to induce the transition as shown in Fig. 2b. For the trajectory in Fig. 2b, the $F_1$ stator angle kept rotated until the simulation ended (Fig. 3b). At the end of the simulation, the $c_{10}$-ring rotated 4×36°, whereas the $F_1$ motor made one transition rotating the chemical states by 120°. The resulting difference, 4×36°-120°=24°, may be absorbed by the $F_1$ stator rotation angle θ (Fig. 3b). The PC1 of the b-subunit distortion changed in parallel with the $F_1$ stator rotation, as in the case of the BC process of the 3-4-3 pathway. In fact, we found many similarities between the AB process which needed four $c_{10}$-ring rotation step (Fig. 3b) and the BC process of the 3-4-3 pathway (Fig. 3c): (i) The $F_1$ stator rotated at the end of the process, (ii) The b-subunit was distorted in parallel with the rotation of the $F_1$ stator, and (iii) four c-ring 36°-rotation steps were necessary to induce a complete transition in $F_1$.

Then, using the 4000th frame snapshot of the AB process trajectory shown in Fig. 2b as the initial structure, we performed BC process simulation. We depict a representative trajectory in Fig. 2d (all the trajectories shown in Supplementary Fig. 1c). αβ1 changed from the DP to the TP state at the beginning of the simulation, as in the trajectory shown in Fig. 2d. Afterward, αβ3 made an irreversible transition from the TP to E states at the ~3100th frame immediately after the third c-ring rotation step in the BC process is complete. Notably, including the AB process, this corresponds to the seventh c-ring rotation step, which is in accordance with the above case. After a while, αβ2 settled down in the post-state, the DP state, at the ~ 4200th frame. This process can be termed the 4-3-3 pathway.

Lastly, we also simulated two other types of CA processes, starting from the 3800th and 4500th frames of the BC process trajectory in Fig. 2d. In both cases, only one trajectory completed the $F_1$ transition by the end of the simulations (Supplementary Fig. 2a, snapshots of 1000 frames from each representative trajectory in Supplementary Fig. 2b). In all cases, the β-subunit was strongly curved from its original position, and αβ3 was found to be dissociated from αβ1 and αβ2, suggesting the failure of these simulations.

#### Supplementary Text 2

##### The ε passing through the side of b-subunit

To complete the $F_1$ structure change in the BC process of the 3-4-3 pathway, the $c_{10}$-ring needs to rotate four 36º-rotation steps, i.e., one step more than in the other processes. The symmetry mismatch between $F_O$ and $F_1$, together with the partial rotation of the $F_1$ stator, primarily explains the need for an extra proton transfer, which will be discussed further in the next section. However, there should be another reason why the BC process, but not the other processes, requires an extra 36º-rotation step. We, therefore, investigated the order of structure transitions among the three αβ pairs within the total 30 trajectories in the AB, BC, and CA processes of the 3-4-3 pathway (Supplementary Fig. 1abd). Of the 30 trajectories, 20 completed all structural transitions in the three αβ pairs. Additionally, we found that in 18 of the 20 cases, the structural change in αβ2 was completed at the end. The structural changes in αβ2 tended to be late in the process. Of the three αβ pairs, αβ2 is unique in that it is in loose contact with the b-subunit, which may cause this delay in the transition.

The trajectory in Supplementary Fig. 5a is a typical trajectory in the BC process (the same as in Fig. 2c). Supplementary Fig. 5b depicts snapshots before and after the structural change in αβ2 (Supplementary Fig. 5a, numbers 1 and 2, respectively). In the pre-state, the β-subunit of αβ2 is trapped by the ε-subunit. At the beginning of the post-state, the ε-subunit pushes the β-subunit outward (moved to the right direction in the figure). Therefore, the timing of the structural change in αβ2 may be controlled by the interactions and positional relationship between αβ2 and the ε-subunit.

To quantitatively evaluate the positional relationship between the α/β-subunits and the ε-subunit, we defined the angles, ∠αOε (and ∠βOε), by the representative position of the $\alpha$-subunit (the $\beta$-subunit), the rotational center, and the representative position of the ε-subunit. The representative positions of the α/β-subunit and the ε-subunit are the mean positions of residues, i.e., the 385th–398th residues and the 101st–112th residues, respectively. The third row of Supplementary Fig. 5a shows the time courses of ∠αOε and ∠βOε of αβ2. We found that ∠βOε changed its sign when αβ2 completed its structural transition to the post-state. The same tendency was observed for the other trajectories in the BC process (Supplementary Fig. 1b).

To make averaging over multiple trajectories possible, we introduced a scaled time. When ∠αOε is 0°, we set the scaled time to 0. In addition, when ∠βOε is 0°, we set the scaled time to 1. Thus, the scaled time is formally defined as:

$$scaled\ time = \frac{original\ time - t_{(\alpha-\varepsilon=0)}}{t_{(\beta-\varepsilon=0)} - t_{(\alpha-\varepsilon=0)}}$$

Using this scaled time for four trajectories in the BC process in which the structural change was correctly completed, we calculated the average χ value of αβ2 along the scaled time (Supplementary Fig. 5c, upper panel). We confirmed that the structural change of αβ2, on average, occurs when the ε-subunit passes through the β-subunit, pushing it outward.

Here, we anticipated that the passage of the ε-subunit through the β-subunit of αβ2, and the resulting transient outward movement of the β-subunit, may be related to the difficulty of structural change in $F_1$ in the BC process. Of the three αβ pairs, only αβ2 had marked contact with the b-subunit, a peripheral stalk. Therefore, the presence of the b-subunit may prevent αβ2 from undergoing transient outward motions necessary for structural changes. To monitor the positional relationship between the b-subunit and the β-subunit of αβ2, Supplementary Fig. 5a (the bottom panel) plots the time course of the angle ∠bOβ defined by the b-subunit, the rotation center, and the β-subunit of αβ2. This angle was defined as positive if the αβ2 β-subunit was located in the counterclockwise direction of the b-subunit (Supplementary Fig. 5b, lower left) and approached zero when the β-subunit was closest to the b-subunit. We see that the b-subunit can touch the β-subunit from the outside in some cases, but not in other cases (Supplementary Fig. 5b, lower left and right, respectively; dashed lines 3 and 4). By plotting the average value of this angle calculated from the four trajectories in the BC process using the scaled time, we found that this angle approaches zero when the χ value of αβ2 changes its sign (Supplementary Fig. 5c lower, Supplementary Fig. 6a). Interestingly, this angle did not cross the zero line in the other six trajectories in which the conformational change was not successful (Supplementary Fig. 6b).

In summary, the β-subunit of αβ2 is supported by the b-subunit from the outside, and thus, it is not easy to move outward. In the BC process of the 3-4-3 pathway, the ε-subunit passes through the side of αβ2, at which the ε-subunit tends to push the β-subunit of αβ2 outward. However, due to its support by the b-subunit from the outside, the β-subunit cannot readily move outward. This may cause an extra bottleneck and contribute to the reason why the BC process requires an extra 36$^{o}$-rotation step of the $c_{10}$-ring.

#### Supplementary Text 3

#### Comparative study of cryo-EM structure models

Our MD simulations showed that ATP synthase exhibits structural changes in multiple elements, which together may resolve the symmetry mismatch. Motivated by these results, in this section, we present a brief comparative survey of structural changes found in the three recent cryo-EM studies: i) the *Bacillus* PS3 ATP synthase by Guo *et al.*, ii) *E. coli* ATP synthase by Sobti *et al.*, and iii) *Polytomella* sp. ATP synthase by Murphy *et al.* Note that all these ATP synthases contain ten c-subunits in the c-ring, making the comparison easy. The first two are from bacteria and resemble each other in all the subunit architecture, whereas the third one is from eukaryote mitochondria and is naturally more divergent from the first two.

To analyze all the structural changes comprehensively, for convenience, we chose the $F_O$ stator, the a-subunit, as the reference point. Relative to this, we calculated the rotation of four parts of the complex around the rotation axis: (i) the rotation of the $c_{10}$-ring, (ii) the rotor distortion defined by the rotation of the upper part of the γ-subunit that interacts with the $F_1$ stator relative to the rotation of the $c_{10}$-ring, (iii) the rotation of the $F_1$ stator relative to the $F_O$ stator, and (iv) the rotation within the $F_1$ motor, defined by the rotation of the γ-subunit relative to the $F_1$ stator $\alpha_3\beta_3$. Note that, by definition, there is a relation: (i)+(ii)=(iii)+(iv). The rotation in (iii) can be attributed to the distortion in the b-subunits. All the rotations are measured from the starting point, which is chosen as the A state of Guo *et al.* or the equivalent one (the rotation state $n = 0$). Then, the B state has an ideal mismatch angle of $3\times36^{o}$ -120 $^{o}$ = -12$^{o}$, whereas the C state has an ideal mismatch angle of $7\times36^{o}$ -240 $^{o}$ = 12$^{o}$.

Results of structure comparisons are given in Table 1.

In the *Bacillus* PS3 ATP synthase structures by Guo *et al*., the B state, which corresponds to the state $n = 3$, has an ideal rotation angle of $c_{10}$-ring, $3\times36^{o}$=108$^{o}$, as well as a nearly ideal $F_1$ motor angle of 119°. The deviation is nearly solely absorbed by the rotation of the $F_1$ stator via the distortion of the b-subunit. In the C state $n = 7$, the $c_{10}$-ring angle deviates from the ideal one by 4$^{o}$, while the $F_1$ stator rotates clockwise by 9$^{o}$. Together, these account for the resolution of the mismatch. The rotor is very rigid with no distortion in both states, and the $F_1$ motor angle deviates slightly from the ideal value.

In the *E. coli* ATP synthase structures by Sobti *et al.*, the 2A (or 2 B), 1A, 1C, and 3A states correspond to the $c_{10}$-ring rotation step $n = 0, 3, 4,$ and $7$. Note that the states $n = 3$ and $n = 4$ possess identical chemical states of $F_1$. Among the four structures, the $F_1$ stator rotates by 15$^{o}$, whereas the rotor was distorted by 8$^{o}$. In addition, the $c_{10}$-ring rotation angles deviate from the ideal angle up to 13$^{o}$. Note that this deviation of the c-ring angle from the ideal angle also serves as an elastic element. These three elements contributed significantly to solving the mismatch. On the other hand, the $F_1$ motor angle changes slightly from the ideal value (~3$^{o}$).

In the *Polytomella* sp., ATP synthase structures by Murphy *et al.* in the 1A, 1F, 2A, 2D, 3A, and 3C states correspond to the $c_{10}$-ring rotation step $n = 0, 1, 3, 4, 7,$ and 8. For simplicity, we chose the two terminal states in each primary step, removing several states in between from the analysis. Among the six structures, the $F_1$ stator rotated by ~28$^{o}$, whereas the rotor was distorted by as much as 22$^{o}$. The $c_{10}$-ring rotation angles deviated from the ideal angle up to 11$^{o}$. In contrast to the above two cases, a marked deviation in the $F_1$ motor angle by 12$^{o}$ was also observed.

Comparing the three cases, we found that the primary source of elasticity used to resolve the symmetry mismatch in these cryo-EM structures was the rotation of the $F_1$ stator relative to the $F_O$ stator, which was realized by the distortion of the b-subunit. However, the distortion of the rotor, as well as the deviation of the $c_{10}$-ring angle from the ideal angle, also contributed to reducing the mismatch. Quantitatively, some differences existed among the three cases, which were primarily attributed to intrinsic differences between different species.

#### Supplementary Text 4

#### The AICG2+ model

As described, we primarily used the AICG2+ model, in which the potential can be expressed as

$$V_{AICG2+} = V_{local} + V_{nonlocal}$$

Here, the local potential has the following terms,

$$V_{local} = \sum_{i \in bond} k_{bond,i}\left(b_i - b_{0,i}\right)^2 + \sum_{i \in angle} V_{angle}(\theta_i) + \sum_{i \in dihedral} V_{dihedral}(\varphi_i)$$

$$+ \sum_{ij\ s.t. j=i+2} \varepsilon_{local\ Go,ij} e^{-(r_{ij}-r_{ij,0})^2/2w_{ij}^2} + \sum_{ij\ s.t. j=i+3} \varepsilon_{local\ Go,ij} e^{-(\varphi_{ij}-\varphi_{ij,0})^2/2w_{\varphi ij}^2}$$

The first term restrains the virtual bond length between adjacent Cα. $b_i$ is the *ith* virtual bond length. The second and third terms are generic statistical potentials for the virtual bond angles and virtual dihedral angles, respectively, which depend on the amino acid type. $\theta_i$ and $\varphi_i$ are the angles between two consecutive virtual bonds and the dihedral angle, respectively. The fourth and fifth terms are structure-based local contact potentials for amino acids i and i+2, and for i and i+3, respectively ( $\varphi_{ii+3}$ represents the dihedral angle defined by the four amino acids, i, i+1, i+2, and i+3). The nonlocal potential is written as:

$$V_{nonlocal} = \sum_{ij \in contact} \varepsilon_{Go,ij}\left[5\left(\frac{r_{ij,0}}{r_{ij}}\right)^{12} - 6\left(\frac{r_{ij,0}}{r_{ij}}\right)^{10}\right] + \sum_{ij \notin contact} \varepsilon_{ex}\left(\frac{d}{r_{ij}}\right)^{12}$$

where the first term is the structure-based contact potential applied to the non-local amino acid pairs that are in contact with the reference structure, whereas the second term is a generic repulsion between non-local amino acid pairs that do not form a contact. $\varepsilon_{Go,ij}$ and $\varepsilon_{local\ Go,ij}$, were determined based on the atomic interaction energy estimate at the reference structure.

The details require further explanation. First, we used the AICG2+ for all the intra-subunit interactions, as well as for all the subunit-subunit pairs that would keep contact throughout the simulation with the default parameter set between the a-subunit and b-subunits, between any c-subunit and γ, between any c-subunit and ε, between α and β that sandwich the catalytic site, between two b-subunits, and between δ and αβ. Second, since the b-subunit, especially b2, had a few missing residues at the C-terminal region, the reference model could not express enough contact with the $\delta$-subunit, and within the AICG2+ model, we set the attractive interaction $\varepsilon_{Go,ij}$ between the b-subunit and $\delta$-subunit to be ten times stronger than the default. Third, for the fragile interface between α- and β-subunits that do not have catalytic sites, we weakened their interaction by 0.7 times, which facilitated the state transitions. Note that the interaction between the b-subunit and αβ is the only excluded volume term without structure-based contact potential.

Lastly, and most importantly, we carefully designed an attractive interaction between the $F_1$ stator α/β-subunit and the $F_1$ rotor $\gamma/\varepsilon$-subunit. Importantly, the rotor $\gamma/\varepsilon$ rotated with respect to the stator $\alpha_3\beta_3$ during the simulation, whereas the reference structure contained native contact information present only in one rotation angle. Based on the set of native contacts for one α- or β-subunit in the reference, we copied them to the same residues in the other two α- or β-subunits. For example, suppose that the $j^{th}$ residue of the $\gamma$-subunit was in contact with the $i^{th}$ residue of the α-subunit 1 in the A state, we added the $j^{th}$ residue of the $\gamma$-subunit and the $i^{th}$ residue of the α-subunits 2 and 3 to the native contact set. When the same residue pair had two types of native contacts originating from different subunits, we chose the native contact with a smaller distance because the latter was considered stronger. Given this duplication of the native contact list, we rescaled $\varepsilon_{Go,ij}$ for the pair between the $\gamma/\varepsilon$ -subunit and the α/β-subunit by 0.5 times. Additionally, we included a sequence-based general hydrophobic interaction between the α/β-subunits and the $\gamma/\varepsilon$ -subunit[1]:

$$V_{HP} = -\sum_i \epsilon_{HP,i} S_i(\boldsymbol{r})$$

where $\epsilon_{HP,i}$ is an experimentally-derived energy scale of transfer free energy for the residue i, and $S_i(\boldsymbol{r})$ is a structure function that approximates buriedness of the residue i.

#### Supplementary Text 5

#### The double basin model for the $F_1$ motor

The double-basin model enables the simulation of the structural change between two reference structures by connecting two potential energies defined by AICG2+[2]. The double-basin potential was defined as the smaller eigenvalue of the following eigenvalue equation:

$$\begin{pmatrix} V_I(R|R_1) & \Delta \\ \Delta & V_I(R|R_2) + \Delta V \end{pmatrix} \begin{pmatrix} c_1 \\ c_2 \end{pmatrix} = V_{MB,I} \begin{pmatrix} c_1 \\ c_2 \end{pmatrix}$$

and, the smaller eigenvalue, the double-basin potential, can be written as follows:

$$V_{MB,I} = \frac{V_I(R|R_1) + V_I(R|R_2) + \Delta V}{2} - \sqrt{\left(\frac{V_I(R|R_1) - V_I(R|R_2) - \Delta V}{2}\right)^2 + \Delta^2}$$

Here, $V_I(R|R_v)$ is the original AICG2+ potential energy for the I-th group in the system, and $v = 1, 2$ represent the pre- and post- reference structures. In our system, we introduced three double basin systems for the three pairs of αβs: αβ1, αβ2, and αβ3. In the AB process simulation, αβ1 should have two basins corresponding to the E and DP states (see Fig. 1c). $V_{\alpha\beta1}(R|R_1)$ contains intra-subunit interactions of α1- and β1-subunits as well as inter-subunit interactions between α1 and β1, and between β1 and α2.

Each double-basin model contains two key parameters $\Delta$ and $\Delta V$. $\Delta$ controls the potential barrier height, and $\Delta V$ represents the relative stability of the two basins. The same value, $\Delta$ was used for each nucleotide state change, independent of the system. In contrast, $\Delta V$ was adjusted for each system and each type of structural change. Because the energy of the three ATP molecules synthesized in one cycle, AB, BC, and CA, was approximately $10 \times 3 = 30\ kcal/mol$, we ensured that the total $\Delta V$ of the whole system was also $+30\ kcal/mol$. The $\Delta$ and $\Delta V$ values for each system in each process are summarized in Supplementary Table 1. In addition, by using the eigenvectors $(c_1, c_2)$, we could define a reaction coordinate that monitored the conformational change χ, defined as

$ln(c_2/c_1)$. When the χ parameter was negative (positive), the system adopted the conformation in pre-state 1 (post-state 2).
